# Combination adjuvants enhance recombinant protein vaccine protection against fungal infection

**DOI:** 10.1101/2021.03.11.434977

**Authors:** Marcel Wuethrich, Hannah E. Dobson, Cleison Ledesma Taira, Uju Joy Okaa, Nikolai Petrovsky, Bruce S. Klein

**Author notes:** Address correspondence to Marcel Wuethrich.

## Abstract

The development of effective vaccines against fungal infections requires the induction of protective, pathogen-specific cell mediated immune responses. Here, we asked whether combination adjuvants based on delta inulin (Advax) formulated with TLR agonists could improve vaccine protection mediated by a fungal recombinant protein, Bl-Eng2, which itself harbors an immunodominant antigen and Dectin-2 agonist/adjuvant. We found that Bl-Eng2 formulated with Advax3 containing TLR9 agonist or Advax8, containing TLR4 agonist, provided the best protection against pulmonary infection with *Blastomyces dermatitidis*, being more effective than Freund’s complete adjuvant or Adjuplex. Advax3 was most efficient in inducing IFN-γ and IL-17 producing antigen-specific T cells that migrated to the lung upon *Blastomyces dermatitidis* infection. Mechanistic studies revealed Bl-Eng2/Advax3 protection was tempered by neutralization of IL-17 and particularly IFN-γ. Likewise, greater numbers of lung-resident T cells producing IFN-γ, IL-17, or IFN-γ^+^ and IL-17^+^ correlated with fewer fungi recovered from lung. Protection was maintained after depletion of CD4^+^ T cells, partially reduced by depletion of CD8^+^ T cells, and completely eliminated after depletion of both CD4^+^ and CD8^+^ T cells. We conclude that Bl-Eng2 formulated with Advax3 is promising for eliciting vaccine-induced antifungal immunity, through a previously uncharacterized mechanism involving CD8^+^ and also CD4^+^ T cells producing IFN-γ and/or IL-17. Although no licensed vaccine exists as yet against any fungal disease, these findings indicate the importance of adjuvant selection for the development of effective fungal vaccines.

**IMPORTANCE:** Fungal disease remains a challenging clinical and public health problem. Despite medical advances, invasive fungal infections have skyrocketed over the last decade and pose a mounting health threat in immune-competent and -deficient hosts with worldwide mortality rates ranking 7th, even ahead of tuberculosis. The development of safe, effective vaccines remains a major hurdle for fungi. Critical barriers to progress include the lack of defined fungal antigens and suitable adjuvants. Our research is significant in identifying adjuvant combinations that elicit optimal vaccine-induced protection when formulated with a recombinant protective antigen and uncovering the mechanistic bases of the underlaying vaccine protection, which will foster the strategic development of anti-fungal vaccines.

## INTRODUCTION

Despite medical advances, invasive fungal infections have skyrocketed over the last decade. They pose a mounting health threat in immune-competent and -deficient hosts with worldwide mortality rates ranking 7^th^, even ahead of tuberculosis(1, 2). The development of safe and effective vaccines remains a major hurdle for fungal prophylaxis. Currently, there are no licensed vaccines against fungi. Subunit vaccines are ideal candidates since they are safe for use in immunocompromised individuals in whom live attenuated vaccines are contraindicated. However, subunit vaccines generally require formulation with a potent adjuvant to be effective. The lack of appropriate adjuvants is one major impediment to developing safe and effective vaccines against fungal pathogens, as is a lack of understanding of the best type of immune response a fungal vaccine would need to elicit to induce maximal protection.

Adjuvants can provide multiple functions, including delivery of antigens to antigen-presenting cells (APCs), formation of antigen depots, chemotaxis, enhanced dendritic cell function and B or T cell co-stimulation. Adjuvants can be combined for enhanced action, such as the combination of innate immune activators such as toll-like receptor ligands with more traditional adjuvant components such as aluminum hydroxide that act as antigen carriers.

Adaptive immunity mediated by CD4^+^ T cells plays the major role in resolution of fungal infections(3, 4), as evidenced by the high incidence of invasive fungal infections in patients with impaired CD4^+^ T-cell immunity. CD4^+^ T cells confer resistance by secretion of T-helper 1 (T_H_1) and T_H_17 cytokines such as IFN-γ, TNF-α, GM-CSF, and IL-17A, which activate neutrophils, monocytes, macrophages and DCs for fungal clearance(4-9). Thus, in order to be protective, adjuvants used for fungal vaccination may need to generate a strong Th1 and Th17 cell immune response.

Advax is based on plant-derived inulin, a polysaccharide that when formulated into delta inulin particles and co-administered with subunit antigens, induces Ag-specific Th1, Th2 and Th17 T cell responses(10-12). Amongst its immunological effects, Advax™ induces a strong chemotactic effect, resulting in the recruitment of leukocytes to the site of vaccination, and engenders vaccine-induced protection against viral and bacterial pathogens(10-18). Advax has not been previously investigated in fungal vaccines. Formulating Advax with TLR agonists enables additional modulation of T cell phenotypes, that may further enhance protective immunity. For example, formulation of the recombinant fusion protein, CysVac containing Ag85B and CysD, with Advax and the TLR9 agonist, CpG, drove an Ag-specific Th1 and Th17 cell response and provided superior protection against aerosol *Mycobacterium tuberculosis* infection(13).

Here, we sought to formulate a recombinant protective fungal antigen, *Blastomyces* endoglucanase 2 (Bl-Eng2), with Advax and various TLR agonists to determine the T helper phenotypes of Ag-specific T cells that best provide protection to fungal infection. We found that Advax-3, which contains the TLR9 agonist CpG, drove a strong balanced Th1 and Th17 cell response and yielded optimal protection against pulmonary infection with the fungus *Blastomyces dermatitidis*. Advax-3 vaccine-induced resistance was mediated by IFN-γ and IL-17 producing CD4^+^ and CD8^+^ T cells.

## RESULTS

### Adjuvant formulations influence anti-fungal T cell development and resistance to fungal infection

To assess the extent to which distinct adjuvant formulations might influence vaccine-induced protection against fungi, we tested nine different adjuvant formulations (Advax 1-9) over multiple experiments (**Table 1**). We compared these Advax formulations to CpG, Freund’s complete adjuvant and Adjuplex™ adjuvant. We formulated the adjuvants with the protective CD4 T cell antigen Bl-Eng2(19), vaccinated mice twice by the subcutaneous (SC) route and challenged them with a pulmonary infection of *B. dermatitidis* two weeks after the boost.

**Table 1:**

| Adjuvant | Formulation |
| --- | --- |
| Advax-1 | inulin polysaccharide adjuvant alone with no TLR agonist |
| Advax-2 | Advax-1 + 10ug TLR9 agonist (CpG55.2) |
| Advax-3 | Aldeltin (special formulation containing alum OH) + 10ug TLR9 agonist (CpG55.2) |
| Advax-4 | Advax-1 + 10ng TLR7/8 agonist |
| Advax-5 | Advax-1 + 200ng TLR4 agonist |
| Advax-6 | Advax-1 + 5ug TLR3 agonist |
| Advax-7 | Advax-1+ 5ug TLR2 agonist |
| Advax-8 | Advax-1 + 1ug TLR4 agonist (5 times more TLR4 agonist than Advax-5) |
| Advax-9 | inulin polysaccharide alone with no TLR agonist- improved antigen binding capacity vs Advax-1 |

At day 7 post-infection, which coincides with the peak influx of primed T cells(20) into the lung, we enumerated the frequencies and numbers of lung cytokine-producing T cells following *ex vivo* stimulation with Bl-Eng2 peptide(19). The tested adjuvant formulations had varying effects on the frequency and cytokine profile of the responding T cells. In the first experiment, the frequencies and numbers of lung IFN-γ and IL-17 producing CD4^+^ T cells post-challenge was significantly increased by the addition of Advax2, 3, 6 and CpG when compared to naïve animals (**Fig. 1A, B, C**). While Advax2 and 6 and CpG induced predominantly single-positive IFN-γ producing or IL-17 producing CD4^+^ T cells, with only a small population of double-positive IFN-γ and IL-17 CD4^+^ T cells (0.49%, 0.03%, 0.35%, respectively), Advax3 was the only adjuvant that induced a relatively large population of double-positive IFN-γ and IL-17 producing cells (3.46%).

**Fig. 1:**
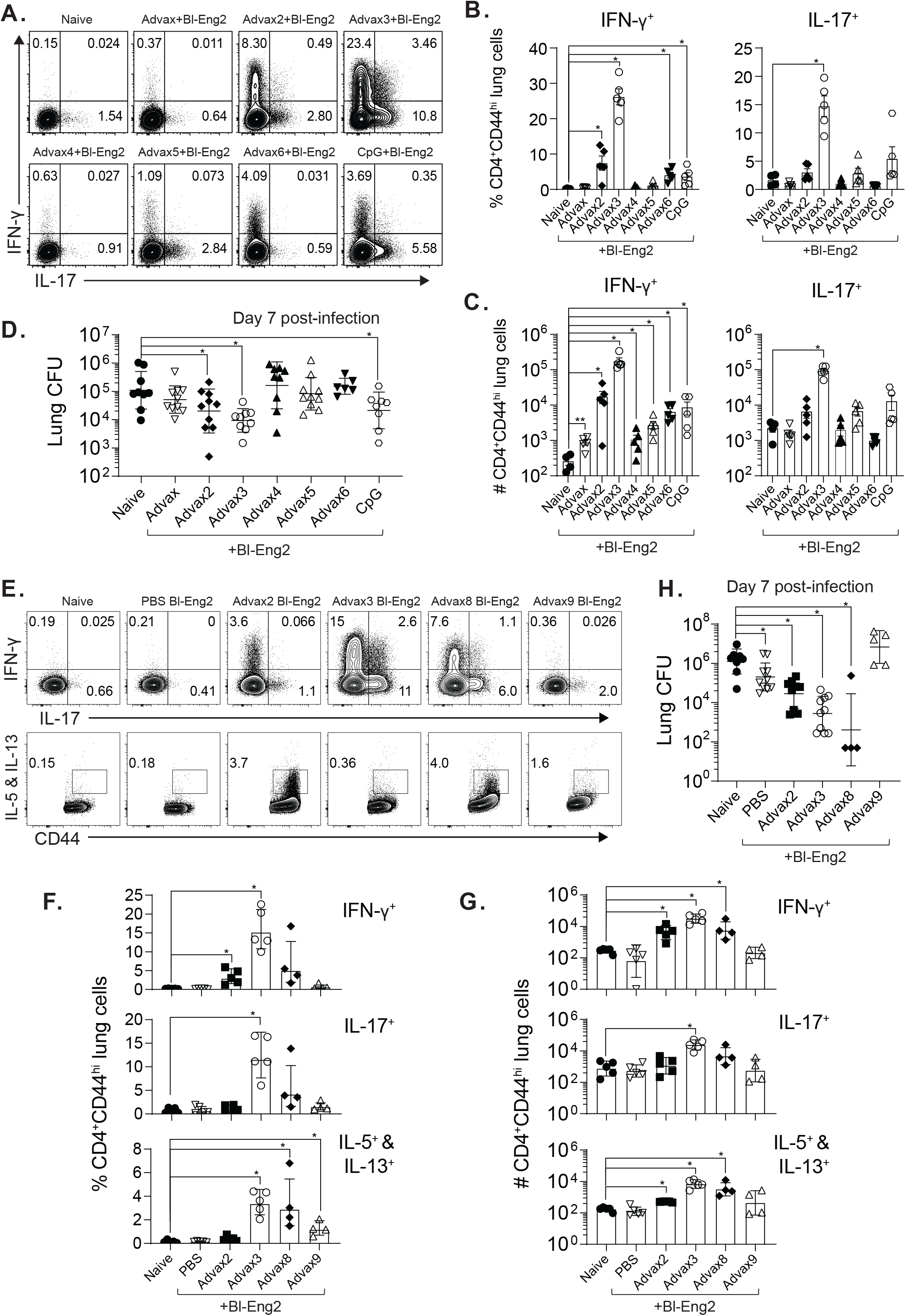
Adjuvant formulations elicit variable Ag-specific T cell responses and protection against fungal infection. **(A)** C57BL6 mice were vaccinated SC with Advax formulations or CpG and 10 µg of soluble Bl-Eng2 protein twice, two weeks apart. Two weeks after the boost, animals were challenged with 2×10^4^ *B. dermatitidis* yeast. Seven days post challenge, mouse lungs were harvested. Dot plots display IFN-γ and IL-17 cytokine production after 5-hours of stimulation with 5 µM of Bl-Eng2 peptide and αCD28. **(B and C)** Corresponding bar graphs show the frequency **(B)** and absolute numbers **(C)** of cytokine-producing CD44^hi^ CD4^+^ T cells (n = 5 mice/group). **(D)** Resistance to infection as determined by lung CFU; graph shows geometric mean with standard deviation (n = 8-10 mice/group). **(E-H)** Repeat experiment with the addition of Advax8 and Advax9. *p < 0.05.

We performed a second experiment to reproduce the initial results with Advax2 and 3, and also investigate two new adjuvant formulations, Advax8 and 9. Advax3 again was associated with the highest numbers of IFN-γ and/or IL-17 secreting CD4^+^ T cells in the lung post-challenge, and the lowest number of IL-5 and IL-13 secreting T cells (**Fig. 1E**), consistent with a balanced enhancement of Th1 and Th17 arms of the adaptive immune response. Advax2, as seen previously, was associated with a significant increase in IFN-γ producing CD4^+^ T cells in the lung post-challenge, consistent with a Th1 bias. Advax-8, like Advax3, was associated with increases in all CD4^+^ T cell populations, although the increase in IL-17 producing CD4^+^ T cells did not reach significance. Advax9 was only associated with an increase in frequency of IL-5 and IL-13 producing CD4^+^ T cells, consistent with an overall Th2 bias. Advax4, 5, 6 and 7, which contained delta inulin plus TLR-7/8, TLR-4, TLR-3 and TLR-2 agonists, respectively, had modest if any effects and were not further pursued.

In the first experiment, formulation of Bl-Eng2 with Advax2, 3 and CpG significantly reduced lung CFU when compared to unvaccinated control mice, with the greatest CFU reduction observed in the Advax3 group (**Fig. 1D**). The level of vaccine protection correlated with the frequencies of single- and double-positive IFN-γ and IL-17 producing CD4 T cells and was highest for Advax3, followed by Advax2 and then CpG **(Fig 1C+F and 3A)**. Notably, those groups that had no increase in double-positive IFN-γ and IL-17 producing CD4^+^ T cells (Advax, 4, 5, 6) also had no significant reduction in lung CFU (**Fig. 1D**).

In the second experiment, formulation of Bl-Eng2 with Advax2, 3 and 8 significantly reduced lung CFU when compared to unvaccinated control mice, with the greatest overall CFU reduction seen with Advax3 and Advax8 (**Fig. 1H**). Interestingly, vaccine formulated with Advax9 showed a polarized Th2 response and no protection, suggesting that a Th2 biased immune response is not beneficial in protecting against lung fungal infection. In summary, Advax3 elicited the highest number of IFN-γ and IL-17 double producing CD4^+^ T cells and reduced lung CFU efficiently when compared to the other Advax and CpG formulations.

### Advax3 compared to Adjuplex and Freund’s adjuvant

We next investigated how Advax3 mediated fungal protection compared to other adjuvants that we, and others, have successfully used to engender subunit vaccine-induced fungal protection. We immunized mice with Bl-Eng2 formulated in Freund’s adjuvant, Adjuplex or Advax3. All three adjuvants activated and expanded IFN-γ-producing Bl-Eng2-specific CD4^+^ T cells. The greatest increase was observed in the Adjuplex group, followed by Advax3 and CFA groups, respectively **(Fig. 2A-C)**. This result was paralleled by the frequency of Bl-Eng2 tetramer-positive CD4^+^ T cells in the lung, which again showed the greatest increase in the Adjuplex group, followed by Advax3 and CFA groups, respectively. However, following infection, Advax3 was associated with the greatest reduction in lung CFU (18-fold) day 4 post-challenge as compared to Adjuplex and CFA adjuvanted vaccines (both 5.6-fold) **Fig. 2D)**.

**Fig. 2:**
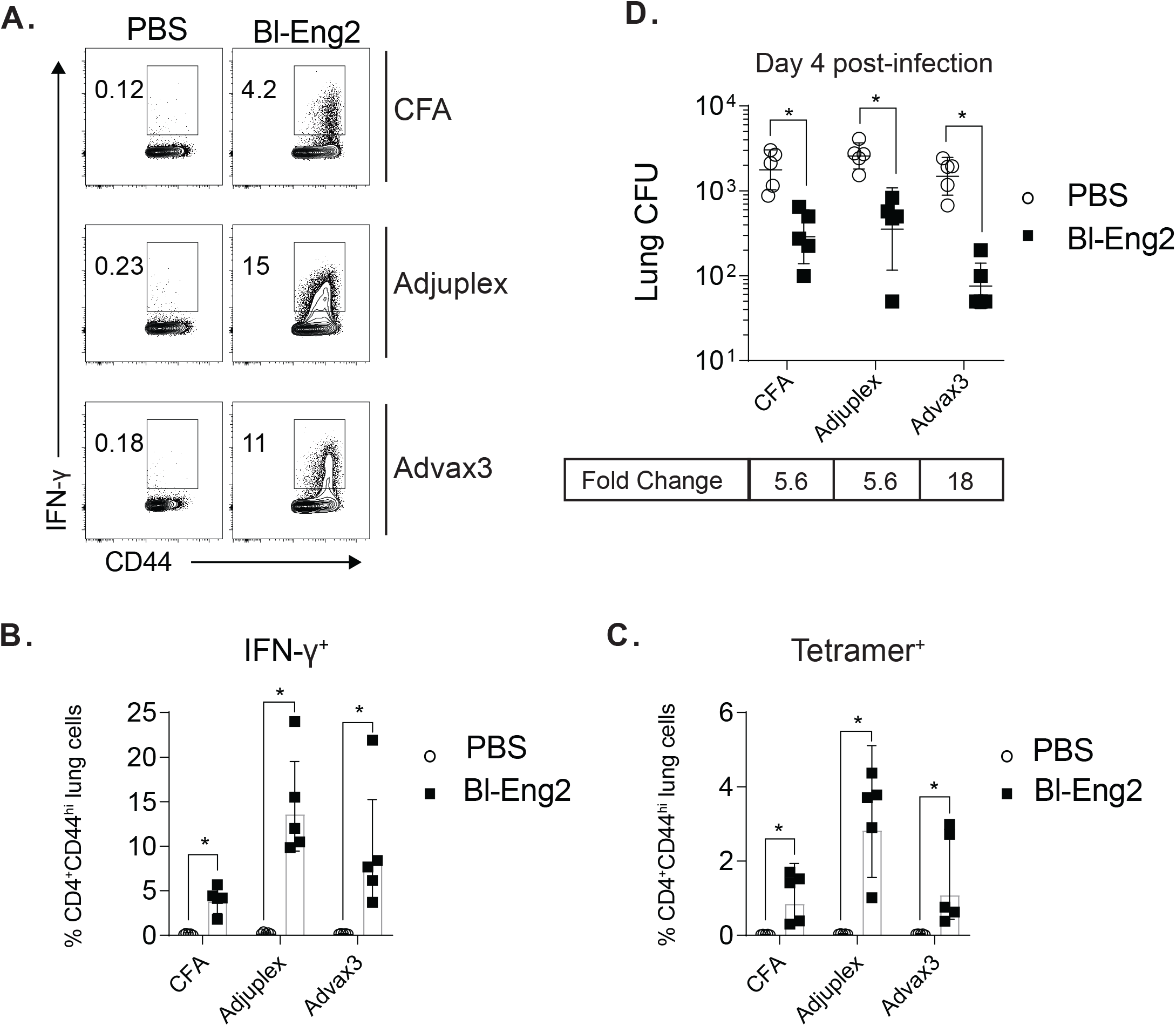
Advax3 provides enhanced fungal protection when compared to Adjuplex and Freund’s adjuvant. C57BL6 mice were vaccinated SC with 10 µg of soluble Bl-Eng2 protein in either Freund’s adjuvant (CFA/IFA), 10% Adjuplex or Advax3 three times, spaced two weeks apart. Two weeks after the last boost, the mice were challenged with 2×10^4^ *B. dermatitidis* yeast. **(A+B)** At day 4 post infection, lung T cells were stimulated with Bl-Eng2 peptide and intracellular IFN-γ was detected by flow cytometry. **(C)** Frequency of Bl-Eng2 peptide tetramer^+^ T cells. **(D)** Lung CFU; graph shows geometric mean with standard deviation and calculated fold change from unvaccinated controls (n = 5 mice/group). *p < 0.05 vs. unvaccinated controls mice.

### Advax3 mediates protection by CD4^+^ and CD8^+^ T cells and IFN-γ and IL-17

We further analyzed the relationship between lung cytokine-producing CD4^+^ T cells and fungal protection. In Bl-Eng2- and Advax3-vaccinated mice, we found a significant, strong correlation between the number of single and double cytokine (IFN-γ and IL-17) producing, Bl-Eng2 specific CD4^+^ T cells in the lung after infection and the reduction in lung CFU **(Fig. 3A)**. We thus sought to investigate whether Advax3-mediated protection is dependent on Th1 and/or Th17 cells. In mice that had been vaccinated with Bl-Eng2 and Advax3, we used anti-cytokine antibodies to neutralize IFN-γ and/or IL-17 prior to challenge and throughout the course of infection. Separately, to further determine which T cell population might play a key role in vaccine protection we used antibodies to deplete CD4^+^ or CD8^+^ T cells alone or together in Bl-Eng2 and Advax3 immunized mice. Two weeks post-infection, when unvaccinated controls were moribund, all the mice were sacrificed and analyzed for weight loss and lung CFU.

**Fig. 3:**
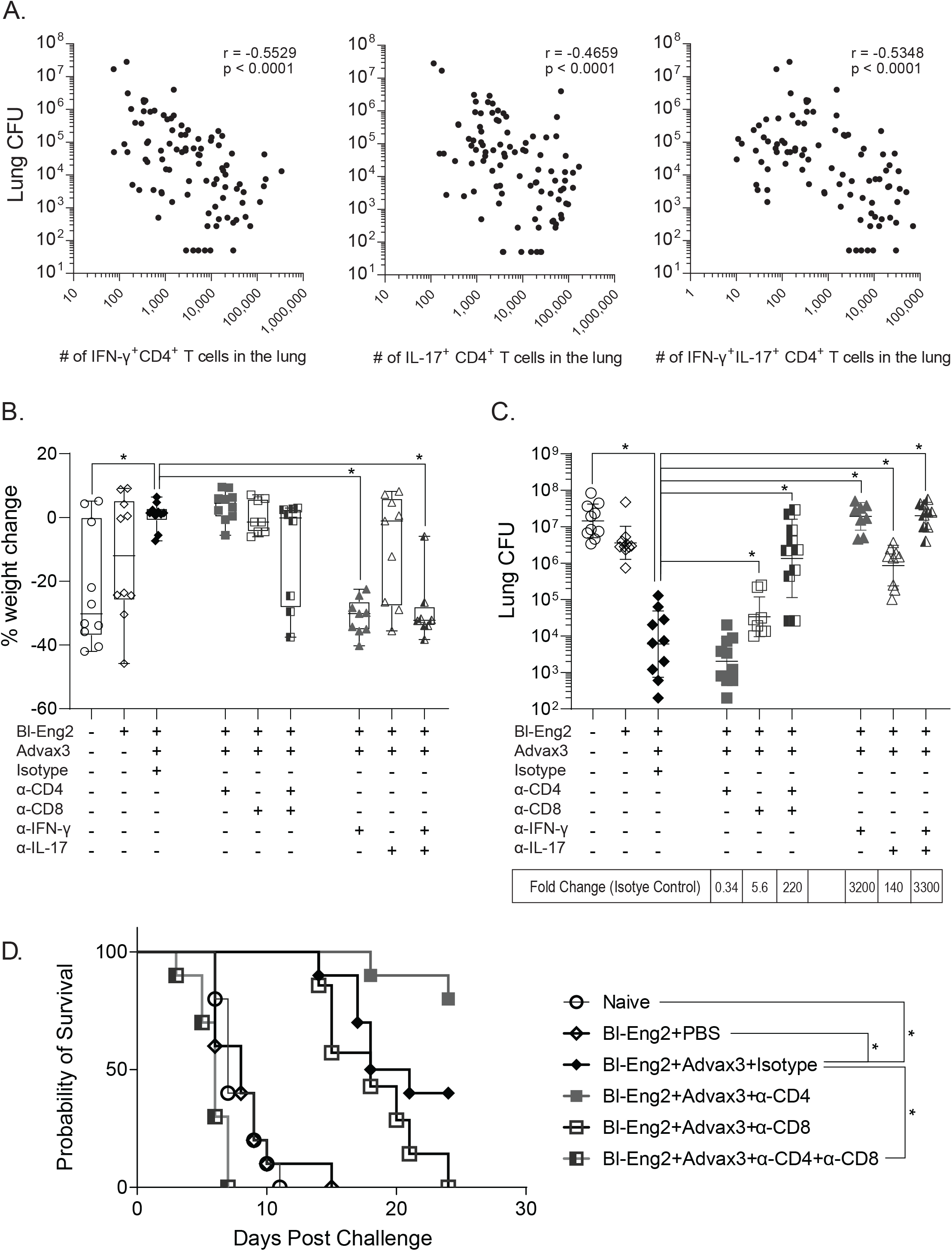
Mechanisms of Advax3-mediated protection. C57BL6 mice were vaccinated SC with 10 µg of soluble Bl-Eng2 protein and Advax3 two times, spaced two weeks apart. Two weeks after the last boost, the mice were depleted of CD4 and/or CD8 T cells or neutralized with anti-IFN-γ or anti-IL-17 mAb and subsequently challenged with 2×10^4^ *B. dermatitidis* yeast. Two weeks post-infection, the mice were sacrificed and analyzed for weight loss **(A)** and lung CFU **(B)**. Graph shows geometric mean with standard deviation and calculated fold change from unvaccinated controls (n = 5 mice/group). *p < 0.05 vs. vaccinated control mice treated with rat IgG. A second cohort of mice was also analyzed for survival for up to 24 days post-challenge **(C)**. *p < 0.05 vs. IgG isotype control treated animals.

Vaccinated mice that were depleted of both CD4^+^ and CD8^+^ T cells or neutralized for IFN-γ or IL-17, or both IFN-γ and IL-17, lost significant amounts of weight and had significantly increased lung CFU compared to Bl-Eng2 and Advax3 vaccinated mice that received isotype control antibody **(Fig. 3B+C)**. Contrary to our expectation, depletion of CD4^+^ T cells alone had little effect on vaccine reduction of lung CFU. By contrast, depletion of CD8^+^ T cells significantly increased lung CFU by 5.6-fold, impairing but not eliminating vaccine protection. Depletion of both CD4^+^ and CD8^+^ T cells abolished protection against weight loss and lung burden of fungi (e.g. CFU), with these double-depleted mice having lung CFU values similar to naive or Bl-Eng2-alone vaccinated controls **(Fig. 3B+C**). Neutralization of IFN-γ resulted in a complete loss of vaccine protection, whereas neutralization of IL-17 had a significant but lesser effect on protection.

We also investigated the impact of CD4^+^ and CD8^+^ T cell depletion on the survival of mice that had been vaccinated with Bl-Eng2 and Advax3 (**Fig 3D**). Naïve controls and animals immunized with Bl-Eng2 antigen alone succumbed to infection by day 14. By contrast, mice immunized with Bl-Eng2 plus Advax3, all survived to day 14 before some became ill, with overall 50% survival at day 24. Depletion of CD8^+^ T cells did not reduce early survival compared to rat IgG control-treated vaccinated mice, but ultimately 100% of CD8^+^ T cell depleted mice succumbed to infection, compared to 60% of the rat IgG control-treated vaccinated mice. Surprisingly, depletion of CD4^+^ T cells increased survival compared to rat IgG treated controls, resulting in over 80% survival at 24 days. Depletion of both CD4^+^ and CD8^+^ T cells resulted in a complete loss of vaccine-induced protection, with all animals in this group succumbing to disease by day 8. Taken together, our results indicate a complex situation where CD8^+^ T cells, but also CD4^+^ T cells, together with IFN-γ and IL-17, all play roles in Advax3-driven vaccine protection. CD4^+^ T cells may potentially function in a double-edged manner, demonstrating the ability to both enhance and diminish vaccine protection.

## DISCUSSION

The hallmark of an effective adjuvant is the ability to elicit the type of adaptive immune response that mediates protective immunity to specific pathogens. In this study, we tested whether Advax inulin-based adjuvant formulations were able to enhance vaccine-induced protection against fungal infection mediated by the protective subunit antigen Bl-Eng2(19). We found that formulating Bl-Eng2 with Advax alone did not augment vaccine protection, whereas the combination of Advax with some TLR agonists but not others did enhance vaccine protection. Combining Bl-Eng2 with Advax3 for vaccination yielded the largest recruitment of Ag-specific Th1 and Th17 cells into the lungs of vaccinated mice following fungal infection. Since vaccine resistance to fungi is principally thought to be mediated by Th1 and Th17 cells(3, 4) we accordingly found that Advax3 which recruited these cells was most efficient in reducing lung CFU, retarding weight loss and prolonging survival of vaccinated mice challenged with *B. dermatitidis*.

Both Advax2 and -3 contain the TLR9 agonist CpG, which yielded a strong induction of IFN-γ producing CD4^+^ T cells. Advax3 induced even greater IFN-γ and IL-17 production by Bl-Eng2-specific T cells compared to Advax2, suggesting that the formulation of Advax3 containing aluminum hydroxide (Table 1) is responsible for the additive effects. In keeping with its ability to increase Th1 and Th17 cell responses, the Advax3 formulation yielded the best vaccine protection. CpG alone combined with Bl-Eng2 also yielded a mixed Th1/Th17 cell response and reduced lung CFU although the combination of Advax with CpG (as in Advax2 and -3) was more effective in driving protective T helper responses and vaccine protection. Advax8 containing a TLR4 agonist also induced a mixed Th1/Th17 cell response and reduced lung CFU efficiently. The TLR4 adjuvant effect was dose dependent as Advax5, which contained five times less TLR4 agonist, showed reduced production of IFN-γ and IL-17 by CD4^+^ T cells and did not reduce lung CFU when compared to Advax8. Formulations containing TLR 2, TLR3 and TLR7/8 agonists were likewise poorly effective at best. Taken together, our data strongly suggest that inclusion of TLR9 (CpG), or possibly TLR4 agonists, with Advax boosts the induction of Ag-specific Th1 and Th17 cells, which are tightly linked with vaccine induced protection against fungi.

Neutralization of IFN-γ completely abolished vaccine-induced protection, whereas neutralization of IL-17 only partially attenuated this protection, indicating the relative importance of IFN-γ to Advax3 mediated vaccine protection. Depletion of both CD4^+^ and CD8^+^ T cells during the expression phase of the vaccine immune response mirrored the phenotypes of IFN-γ and IL-17 neutralization, suggesting that IFN-γ and IL-17 produced by T cells mediate Advax3 vaccine resistance. Depletion of CD8^+^ T cells increased lung CFU and reduced survival compared to isotype control antibody-treated mice, whereas depletion of CD4^+^ T cells did not reverse the vaccine effect on lung CFU and actually improved overall survival. These results indicate that CD8^+^ T cell subsets are the primary contributor to Advax3 vaccine immunity, but with some CD4^+^ T cells still able to compensate for and provide protection in the absence of CD8^+^ T cells. Notably, subsets of CD4^+^ T cells are able to produce IFN-γ and IL-17 and it is likely these subsets that are able to promote protection in the absence of CD8^+^ T cells. Nevertheless, the data indicate that another subset of CD4^+^ T cells contributes to disease pathogenesis given the increased survival of vaccinated mice after CD4^+^ T cell depletion. As adjuvant groups that mounted solely Th2 cytokine responses showed minimal protection (e.g. Advax 9), we speculate that CD4^+^ T cells may have a dual role in vaccine protection, with Th1 and Th17 subsets contributing to protection via IFN-γ and IL-17 production and the Th2 subset contributing to disease progression via known inhibitory effects of Th2 cytokines such as IL4 and IL10 on IFN-γ and IL-17 production.

The strong contribution of CD8^+^ T cells to protection conferred by Advax3 was unexpected, as we have previously observed a primary role for CD4^+^ T cells in mediating vaccine resistance. Those results were observed when we immunized mice with a live attenuated strain of *B. dermatitidis* or with subunit vaccines containing the conserved antigens Calnexin or Bl-Eng2 together with other adjuvants(19, 21, 22). In fact, we reported a clear hierarchy in priming of CD4^+^ T cells over CD8^+^ T cells when using the live attenuated strain to induce vaccine immunity (21, 23). In a one study, we noted that depletion of CD4^+^ T cells, but not CD8^+^ T cells, during the vaccine effector phase abolished vaccine protection (21). In a second study, we reported differential requirements by CD4^+^ and CD8^+^ T cells for co-stimulation by the CD40/CD40L axis during the vaccine priming phase(23). CD4^+^ T cells, but not CD8^+^ T cells, required CD40/CD40L co-stimulation. Most importantly, antifungal CD8^+^ T cells failed to become activated and mediate resistance in CD40^-/-^ and CD40L^-/-^ mice when CD4^+^ T cells were present, indicating that the presence of CD4^+^ T cells impeded priming of CD8^+^ T cells (23). Thus, the prominent role of CD8^+^ T cells over CD4^+^ T cells in mediating Advax3 induced vaccine immunity was unanticipated and merits further investigation. Our results raise the possibility that the Advax3 adjuvant was key to this observation, due to its ability to strongly prime CD8^+^ T cells, thereby preventing normal CD4^+^ T cell suppression. Consistent with our findings in the present study, Advax 2 and 3 have both been shown to induce cross presentation to CD8^+^ T cells and induce high levels of *in vivo* CTL killing of CD8 peptide-labelled target cells (11, 24). Our findings on the apparent ability of Advax3 to directly engage CD8^+^ T cells in mediating antifungal vaccine protection even in the absence of CD4^+^ T cells could have important implications for successful immunization of patients with compromised CD4^+^ T cell immunity who are vulnerable to fungal infections.

## METHODS

### Fungi

Wild-type, virulent *B. dermatitidis* ATCC strain 26199 was used for this study and grown as yeast on Middlebrook 7H10 agar with oleic acid-albumin complex (Sigma) at □ LJC.

### Mouse strains

Inbred wild type C57BL/6 mice obtained from Jackson Laboratories were bred at our facility. Male and female mice were 7-8 weeks old at the time of these experiments. Mice were housed and cared for as per guidelines of the University of Wisconsin Animal Care Committee who approved all aspects of this work.

### Vaccination and infection

For Advax comparisons, 10 µL of Advax adjuvant (Advax - Advax9) was combined individually with 10 µg of soluble Bl-Eng2 protein and diluted to 200 µL with PBS. The vaccine was delivered in this 200 µL dose by subcutaneous injection. Mice received one boost two weeks following the initial vaccination.

In adjuvant comparison experiments, Bl-Eng2 was formulated in CFA/IFA, 10% Adjuplex and Advax3. 50 µL of CFA was combined with 10 µg of soluble Bl-Eng2 protein and diluted to 200 µL with PBS. CFA vaccines were then sonicated and injected subcutaneously in 200 µL doses. Adjuplex vaccines were also delivered in 200 µL doses by adding 10 µg of soluble Bl-Eng2 protein to 20 µL of Adjuplex adjuvant and diluting to 200 µL with PBS. Advax3 was formulated as described above. Following the initial vaccination, mice received two subsequent boosts with IFA, 10% Adjuplex and Advax3, spaced two weeks apart.

Vaccinated animals were infected with *B. dermatitidis* yeast two weeks after the last boost. The mice received 2×10^4^ yeast by aspiration as described(19). Mice were euthanized for cellular and CFU analysis either at day 4 post-infection or when they first appeared moribund.

### T cell stimulation and flow cytometry

The lungs were dissociated in Miltenyi MACS tubes and digested with collagenase D (1mg/ml) or collagenase B (2mg/ml, for Influenza) and DNase (1ug/ml) for 25 minutes at 37°C. The digested lungs were resuspended in 5ml of 40% percoll (GE healthcare, cat 17-0891-01); 3 ml of 66% percoll was underlaid. Samples were spun for 20 minutes at 2,000 rpm at room temperature. The lymphocytes layer was collected and resuspended in complete RPMI (10% FBS, and 100ug/ml Penicillin/Streptomycin). For the influenza vaccine model, the percoll separation was not performed. Instead after digestion of the lungs, the red blood cells were lysed with ACK buffer, the samples filtered in 40 µM cell strainers and the cells resuspended in complete RPMI. For *ex vivo* stimulation, lung T cells were incubated at 37°C for 5 hours with 5 µM Bl-Eng2 peptide and 1µg antimouse CD28 (BD, cat 553294). After 1h, GolgiStop (BD, cat 554724) was added to each well. All FACS samples were stained with LIVE/DEAD Fixable Near-IR Dead Cell Stain (Invitrogen) and Fc Block (BD) for 10 minutes at room temperature. T cells were stained with Bl-Eng2 tetramer for 1 hour at room temperature. Then, the cells were stained for surface antigens and intracellular cytokines. All panels included a dump channel (B220, CD11b, CD11c and NK1.1). 50 µl of AccuCheck Counting Beads (Invitrogen, cat PCB100) was added to the samples to determine the absolute cell count. Samples were acquired on a LSR Fortessa at the University of Wisconsin Carbone Cancer Center Flow Lab.

### Surface panel cocktail for Bl-Eng2 peptide experiments

CD45 AF488 (Biolegend, clone 30-F11, cat 103122), MHC class II tetramer-PE, CD4 BUV395 (BD, clone GK1.1, cat 563790), CD8 PerCP-Cy5.5 (Biolegend, clone 53-6.7, cat 100734), CD44 BV650 (Biolegend, clone IM7, cat 1033049), CD11b APC (Biolegend, clone M1/70, cat 101212), CD11c APC (Biolegend, clone N418, cat 117310), NK1.1 APC (Biolegend, clone PK136, cat 108710), B220 APC (Biolegend, clone RA3-62B, cat 103212).

### *In vivo* T cell depletion and cytokine neutralization

Vaccinated mice were depleted of T cell subsets by IV injection of 100 μg anti-CD4 (clone GK1.5) and/or anti-CD8 (clone 53-6.7) or rat IgG as a negative control two hours before challenge and weekly thereafter. Cytokines were neutralized IV injection of 100 μg anti-IFN-γ (clone XMG1.2) and/or IL-17A (clone 17F3) mAbs two hours before challenge and every other day thereafter. All antibodies were purchased from BioXcell, Lebanon, NH, USA.

### Statistics

All statistics for flow cytometry and CFU were calculated in Prism (GraphPad). An unpaired two-tailed t test was used to calculate significance between naïve and vaccinated groups in either absolute number or percentage of cells. For comparisons of absolute cell counts, data were log transformed before t-tests were conducted. CFU data were also log transformed and an unpaired two-tailed t test with Welch’s correction was used to calculate p values. Survival outcomes were analyzed in Prism (GraphPad) using the Bonferoni method adjusting for multiple comparisons. A p≤ 0.05 was considered statistically significant.

## Ethics statement

The animal studies performed were governed by protocols M005891 as approved by the IACUC committees of the University of Wisconsin-Madison Medical School. Animal studies were compliant with all applicable provisions established by the Animal Welfare Act and the Public Health Services (PHS) Policy on the Humane Care and Use of Laboratory Animals.

## ACKNOWLEDGEMENTS

The work was supported by National Institutes of Health grants AI093553 (MW), AI035681 (BK), AI040996 (BK). Flow samples were collected at the University of Wisconsin Carbone Center Flow Lab on a BD LSR Fortessa that was purchased with the NIH shared instrumentation grant 1S100OD018202-01. NP and development of Advax adjuvant was supported by funding from National Institute of Allergy and Infectious Diseases of the National Institutes of Health under Contracts HHSN272201800044C, HHS-N272201400053C, HHS-N272200800039C and U01-AI061142.

